# Estimating alpha, beta, and gamma diversity through deep learning

**DOI:** 10.1101/2022.01.12.475997

**Authors:** Tobias Andermann, Alexandre Antonelli, Russell L. Barrett, Daniele Silvestro

**Affiliations:** Department of Biological and Environmental Sciences, University of Gothenburg, Sweden; Gothenburg Global Biodiversity Centre, University of Gothenburg, Sweden; Department of Biology, University of Fribourg, Switzerland; Swiss Institute of Bioinformatics, Fribourg, Switzerland; Department of Plant Sciences, University of Oxford, United Kingdom; Royal Botanic Gardens, Kew, Richmond, Surrey, United Kingdom; Royal Botanic Gardens, Sydney, New South Wales, Australia; School of Biological Sciences, The University of Western Australia, Crawley, Australia

**Author notes:** **Correspondence:** Tobias Andermann.

## Abstract

The reliable mapping of species richness is a crucial step for the identification of areas of high conservation priority, alongside other value considerations. This is commonly done by overlapping range maps of individual species, which requires dense availability of occurrence data or relies on assumptions about the presence of species in unsampled areas deemed suitable by environmental niche models. Here we present a deep learning approach that directly estimates species richness, skipping the step of estimating individual species ranges. We train a neural network model based on species lists from inventory plots, which provide ground truthing for supervised machine learning. The model learns to predict species richness based on spatially associated variables, including climatic and geographic predictors, as well as counts of available species records from online databases. We assess the empirical utility of our approach by producing independently verifiable maps of alpha, beta and gamma plant diversity at high spatial resolutions for Australia, a continent with highly contrasting diversity patterns. Our deep learning framework provides a powerful and flexible new approach for estimating biodiversity patterns.

## 1 Introduction

Since the very beginnings of biogeographic research, the estimation and extrapolation of species diversity has been of foremost interest (Humboldt, 1817; Arrhenius, 1921). It is well established that species diversity is distributed unevenly across space, generally following a latitudinal gradient, with increasing diversity from the poles toward the equator (MacArthur, 1965). On a regional level, it has been found that there are substantial differences in species richness among habitats, such as between a forested area and an open grassland (MacArthur, 1965). These observed spatial patterns have led to the formulation of three levels of species diversity: alpha, beta, and gamma diversity (Whittaker, 1960).

Alpha diversity (Whittaker, 1960) refers to diversity on a local scale, describing the species diversity (richness) within a functional community. For example, alpha diversity describes the observed species diversity within a defined plot or within a defined ecological unit, such as a pond, a field, or a patch of forest. The scale of such ecological units depends on the organism group of interest; while for birds a defined forest or grassland transect of several hundred m^2^ to several km^2^ may be appropriate to describe a species community, for insects this could be a single tree. For plants, alpha diversity is often equated to the count of species identified during the inventory of a vegetation plot of defined size (Revermann et al., 2016).

Beta diversity, on the other hand, describes the amount of differentiation between species communities (Whittaker, 1960). Unlike the other levels of species diversity, the exact interpretation and quantification of beta diversity varies significantly across studies (see Tuomisto, 2010a, 2010b for a detailed review on this topic). Originally, beta diversity was defined as the ratio between gamma and alpha diversity (*β* = *γ*/*α*, sensu Whittaker, 1972). Today, one of the more commonly used measures of beta diversity is the Sørensen dissimilarity index (see Methods below for more detail), which captures spatial turnover as well as differences in diversity between sites (Koleff et al., 2003).

Gamma diversity describes the overall species diversity across communities within a larger geographic area (Whittaker, 1960). It is often summarized across biogeographic or political units, such as ecoregions or countries (Kier et al., 2005; Brummitt et al., 2021). Alternatively, studies commonly summarize gamma diversity within cells of a spatial grid of fixed cell-size (Goldie et al., 2010; Thornhill et al., 2016). While alpha diversity represents the actual species diversity that can be measured at a given site, gamma diversity more broadly and loosely describes the diversity of species that can be found in the whole area. Gamma diversity is the most communicated level of species diversity when referring to biodiversity hotspots, with tropical regions, in particular the Neotropics, showing the globally highest gamma diversity values (Raven et al., 2020). Alpha diversity, on the other hand, shows different areas of maximum diversity, dependent on the size of the area surveyed, with temperate grasslands showing among the highest species richness on small plots (Wilson et al., 2012).

While alpha diversity can be directly counted for small plot sizes, for example during species inventories, this requires much effort and thus cannot be scaled up to large areas or whole continents. Therefore, many studies apply some form of modeling and estimation to derive diversity maps for larger areas. For example, gamma diversity is often inferred by modelling individual species distributions and adding these up to derive the total number of species that occur in a given area (Mutke and Barthlott, 2005; Barthlott et al., 2007). However, this approach has been shown to introduce substantial errors, when cross-checking the diversity predictions with actual species counts in selected grid cells (Aranda and Lobo, 2011). A general shortcoming of these methods is that usually there is not sufficient data available to reliably model the individual ranges for each species. This problem intensifies with the size of the target group for which to estimate diversity patterns. In some cases, total species diversity is extrapolated for larger groups, based on a selected subset of taxa with good data coverage, under the simplistic assumption that the diversity patterns revealed by these taxa are representative for others (Kier et al., 2005), which is however often not the case (Ritter et al., 2019).

Alternative approaches have been applied to the task of diversity estimation and mapping, which skip the step of modelling individual species ranges. These often involve using occurrence records, floras, and checklists for large biogeographic regions (Mutke and Barthlott, 2005; Kreft and Jetz, 2007). While such approaches do not require to model distributions of individual species, they are particularly vulnerable to biases in data collection, as some taxa may be better represented in some checklists and biodiversity repositories than others. When models are applied, they usually assume a single diversity value within each of the regions analyzed, without accounting for fine-scale local fluctuations within these (sometimes large) areas. Although it may be possible to interpolate diversity values to a finer resolution using spatial autocorrelation of associated variables such as climatic predictors (Kreft and Jetz, 2007), such gap filling may be difficult to verify and often provides a false sense of confidence for data-poor regions.

With the increasing availability of continental and global vegetation plot databases (Chytrý et al., 2016; Bruelheide et al., 2019; Sabatini et al., 2021), a new data source with extended spatial coverage has become widely available for the task of large-scale diversity estimation. Recently, Večeřa et al. (2019) showed the potential of machine learning models (random forest models) to estimate the expected diversity for fixed size vegetation plots (alpha diversity), based on climatic and other predictors, when trained on alpha diversity data from vegetation plot databases. However, to our knowledge, available machine learning models cannot extrapolate vegetation plot data to larger areas and do not provide estimates of multiple metrics of biodiversity.

Here we present a deep learning framework that uses neural network models (deep learning) to predict alpha, beta, and gamma diversity. Our approach requires neither geographic data of individual species, nor the manual extrapolation of species richness using methods such as species–area curves (Kier et al., 2005). Instead, our models inherently learn the species-area relationships, allowing prediction of the three diversity metrics at user-defined spatial scales. The models learn to predict plant diversity based on climatic and geographic predictors, measures of human impact, and sampling effort.

We selected plot-based vegetation survey data from Australia (vascular plants; Tracheophyta) to empirically test the effectiveness of these neural network models to predict diversity patterns and validate our methodology. Australia, as an island continent, has the advantage of a clear delimitation of natural boundaries; it has high natural diversity and uneven biological sampling (González-Orozco et al., 2014; Cook et al., 2015; Laffan et al., 2016); high spatial heterogeneity with well-defined and contrasting biomes (Byrne et al., 2008, 2011; Macintyre and Mucina, 2021); a relatively well-documented vascular flora with reliable national databases (https://avh.chah.org.au/; Sparrow et al., 2021) that feed into the Global Biodiversity Information Facility (GBIF, gbif.org); good climatic data (http://www.bom.gov.au/climate/data/); and a large number of freely available plot-based vegetation records suitable for training deep learning frameworks (Sabatini et al., 2021).

## 2 Methods

### 2.1 Vegetation plot data

The values of alpha, beta, and gamma diversity used in this study to train the neural network models were derived from vegetation plot data (species inventories). We downloaded these data from the sPlotOpen database (Sabatini et al., 2021), only using plots where all vascular plants had been assessed. This resulted in a total of 7,896 vegetation plots for Australia (Fig. 1). For each vegetation plot, we compiled its area (in m^2^) and the list of species identified. From each of these sites we compiled measures for alpha, beta, and gamma diversity as described in more detail below (Fig. 1), which we used to train our models.

**Figure 1:**
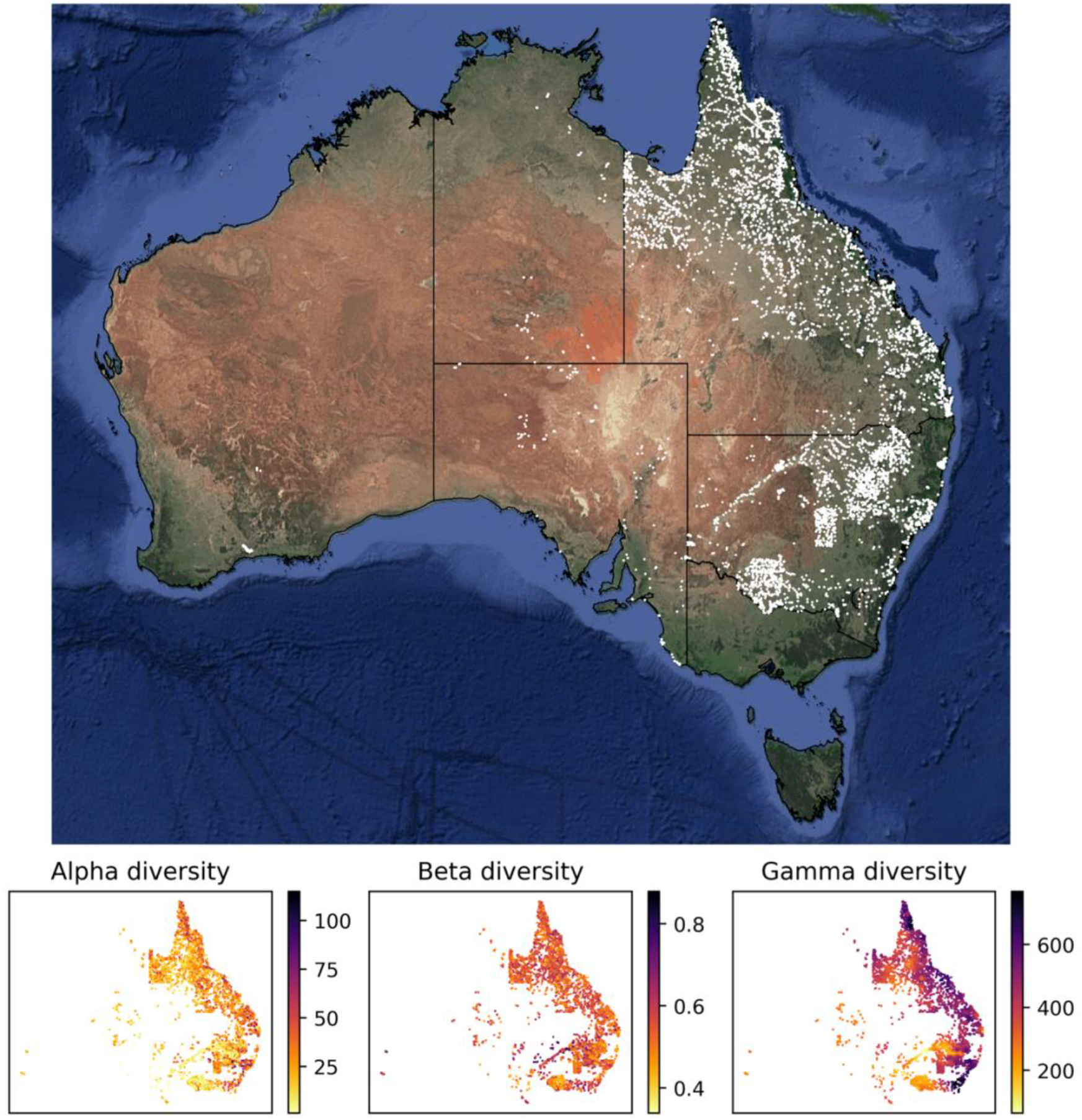
Sites with vegetation plot data used in this study for model training and evaluation. Most of the vegetation plot sites used in this study (white points, 7,896 sites) are located in the easternmost two Australian states Queensland (northeast) and New South Wales (center east). Our uncertainty quantification (Fig. 4) addresses these spatial biases in the underlying data, showing higher prediction uncertainty in areas with low data coverage. The panels below the map show the compiled measure of alpha, beta, and gamma diversity for all vegetation plot sites. The satellite image of Australia was downloaded via ggmap (Kahle and Wickham, 2013).

Calculating gamma diversity required the definition of a surrounding area, preferably containing other vegetation plots, to determine the overall diversity found within the cumulative species lists of several neighboring vegetation plots (Fig. 2). To ensure that the same number of vegetation plots was used for calculating the gamma diversity of each site, we defined as the surrounding area a circle around each site encompassing exactly the N nearest neighbors (vegetation plots). The gamma diversity for each site was then determined as the number of unique species extracted from the species lists of the N nearest neighbors within the encompassing circle. After compiling diversity estimated for different values of N (Supplementary Figs. S1-S7), we chose an N of 50 for all models in this study, as this value led to the best compromise between a visually discernible spatial structure in the resulting beta and gamma diversity values, while also highlighting regional heterogeneity (Supplementary Fig. S3).

**Figure 2:**
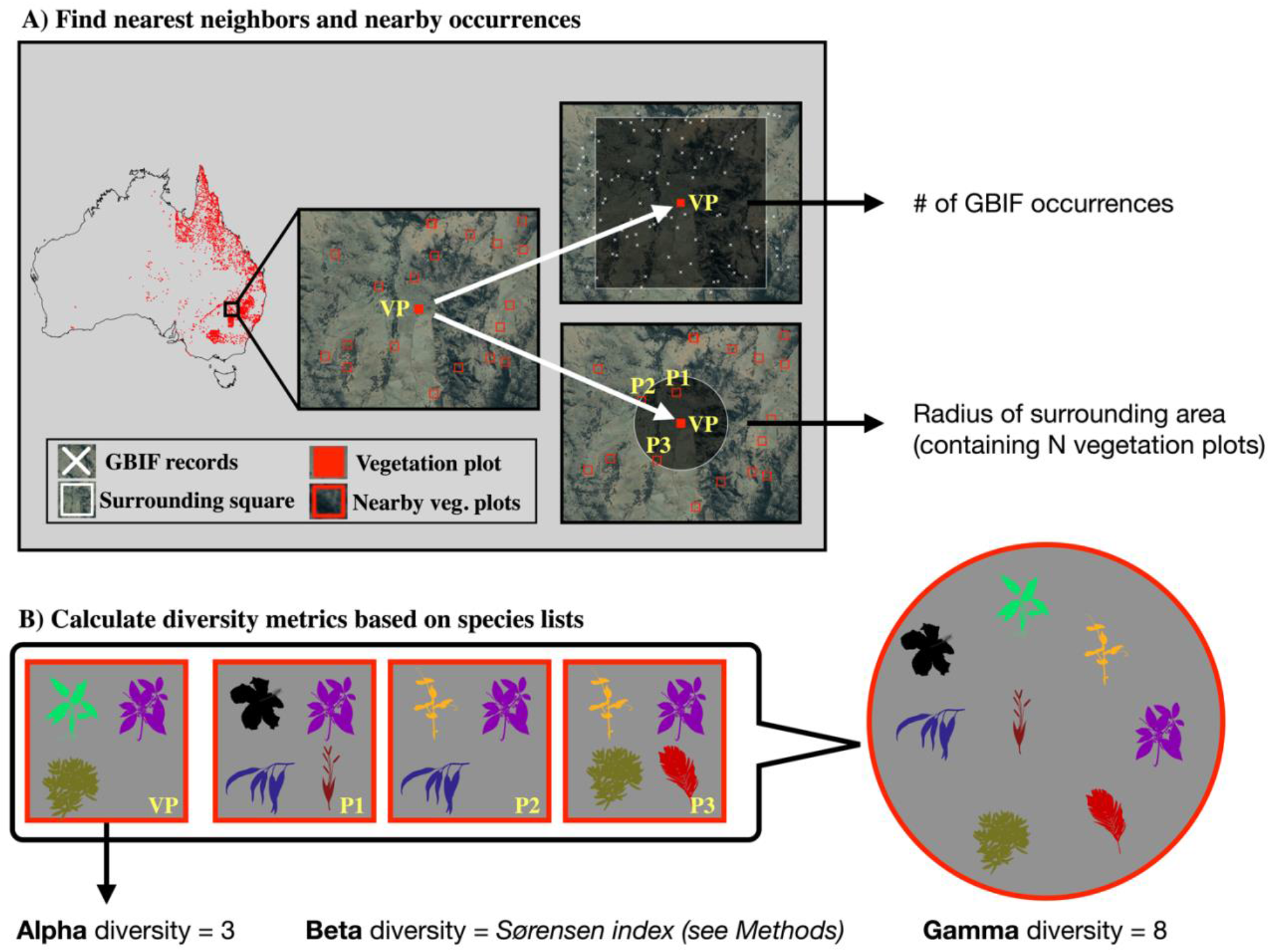
Calculation of diversity measures from vegetation plot data. For a given vegetation plot (VP, solid red square, panel A) we identified the N nearest neighboring vegetation plots in space (N=3 in this example, represented by plots P1-P3). We exported the radius of the smallest circle encompassing all N neighbors as a feature for model training. Additionally, we exported the number of GBIF occurrences within a square of 10×10 km size around the given vegetation plot, as a measure of sampling effort in the general area. Having identified the nearest neighbors (P1-P3), we compared the species lists of these vegetation plots with the focal vegetation plot (VP, panel B). Alpha diversity represents the number of species found in the focal vegetation plot (VP), while gamma diversity represents the total diversity consisting of all species identified among the focal and neighboring vegetation plots. Beta diversity was calculated using the multiple-site Sørensen dissimilarity index (see Methods), based on the differences in species composition found among the selected vegetation plots.

The radius of this encompassing circle varied between sites, depending on the proximity of other vegetation plots relative to the given site. This radius was used as a feature in our models, allowing the neural network to learn the expected associations between gamma diversity and the size of the area for which it was calculated (the species-area curve), which we used later when making predictions with this model to adjust the spatial resolution of the predictions.

Finally, beta diversity was calculated using the multiple-site implementation of the Sørensen dissimilarity index (*β_sor_*), following the definition in (Baselga, 2010). For a given focal site *j* with N neighbors, we defined the focal site index as *j*=*N*+*1*. We iterated through the N neighboring sites (*i*) and applied the formula:

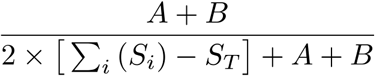

with

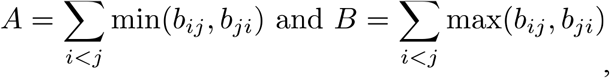

where *b_ij_* and *b_ji_* are the number of species only present in site *i* and site *j*, respectively, *S_i_* is the total number of species in site *i* (alpha diversity from vegetation plot), and *S_T_* is the total number of species in all sites combined (gamma diversity).

### 2.2 Feature generation

The alpha, beta, and gamma diversity metrics described above were used as labels to train three models based on a range of different features, one for each diversity metric. To ensure approximately equal size of all grid cells for the raster-based data used in this study, we transformed all spatial data into the cylindrical equal-area (CEA) projection, centered at 30 degrees latitude south of the equator.

As a general measure of sampling effort, we compiled the number of recorded species occurrences, available on GBIF which were found in the vicinity of a given site. We first downloaded all non-fossil vascular plant (Tracheophyta) occurrences for Australia from GBIF based on human observations and not flagged for geospatial issues (https://doi.org/10.15468/dl.kbq3d7). This includes both native and naturalized species, the latter having uneven spatial distributions related to broad disturbance histories in Australia (Leishman et al., 2017). This resulted in 13,580,191 occurrence records. We then discarded any records with non-binomial species names and cross-checked names of the remaining records against the World Checklist of Vascular Plants, a continuously updated collection of reviewed plant species names (Govaerts et al., 2021). This resulted in 12,622,786 remaining GBIF records. For each site, we defined a 10×10 km window centered on the site’s coordinates; we then counted all GBIF occurrences within this window as a measure of sampling effort (Supplementary Fig. S8), as well as the number of species found in the GBIF records as a diversity proxy. Both counts were used as individual features in our models.

We also compiled climatic and anthropogenic features for each site. First, we downloaded raster data for 19 bioclimatic variables (BIO1-BIO19) as well as data on elevation from the WorldClim database (worldclim.org, Fick and Hijmans, 2017). Second, we downloaded raster data on human footprint from wcshumanfootprint.org (Venter et al., 2016), which reflects the magnitude of human disturbance, including information on human population density, agricultural land use, presence of roads and several other data sources. There is a high coincidence between population density, agricultural development, and high biodiversity regions in Australia (Keith and Auld, 2017). All data rasters were downloaded at a resolution of 0.5 minutes of a degree (~1×1 km grid). The complete list of features (n=27) extracted for each site is shown in Table 1. All feature values were rescaled to range between 0 and 1 before being used as input in the neural network.

**Table 1:** Features used in the NN models.

| Index | Feature name | Data source | Selected 27 | Selected 8 | Selected 6 |
| --- | --- | --- | --- | --- | --- |
| 1 | Longitude | sPlotOpen | X | X |  |
| 2 | Latitude | sPlotOpen | X | X |  |
| 3 | Sampling effort | gbif.org | X |  |  |
| 4 | # of detected species | gbif.org | X |  |  |
| 5 | Human footprint | wcshumanfootprint.org | X | X | X |
| 6 | Elevation | WorldClim | X | X | X |
| 7 | BIO1 (Annual mean temperature) | WorldClim | X | X | X |
| 8 | BIO2 (Mean diurnal range) | WorldClim | X |  |  |
| 9 | BIO3 (Isothermality) | WorldClim | X |  |  |
| 10 | BIO4 (Temperature seasonality) | WorldClim | X |  |  |
| 11 | BIO5 (Max. temp. warmest month) | WorldClim | X |  |  |
| 12 | BIO6 (Min temp coldest month) | WorldClim | X |  |  |
| 13 | BIO7 (Temperature annual range) | WorldClim | X |  |  |
| 14 | BIO8 (Mean temp. wettest quarter) | WorldClim | X |  |  |
| 15 | BIO9 (Mean temp. driest quarter) | WorldClim | X |  |  |
| 16 | BIO10 (Mean temp. warmest quarter) | WorldClim | X |  |  |
| 17 | BIO11 (Mean temp. coldest quarter) | WorldClim | X |  |  |
| 18 | BIO12 (Annual precipitation) | WorldClim | X | X | X |
| 19 | BIO13 (Precipitation wettest month) | WorldClim | X |  |  |
| 20 | BIO14 (Precipitation driest month) | WorldClim | X |  |  |
| 21 | BIO15 (Precipitation seasonality) | WorldClim | X |  |  |
| 22 | BIO16 (Precipitation wettest quarter) | WorldClim | X |  |  |
| 23 | BIO17 (Precipitation driest quarter) | WorldClim | X |  |  |
| 24 | BIO18 (Precipitation warmest quarter) | WorldClim | X |  |  |
| 25 | BIO19 (Precipitation coldest quarter) | WorldClim | X |  |  |
| 26 | Vegetation plot size | Based on sPlotOpen data | X | X | X |
| 27 | Neighborhood radius | Based on sPlotOpen data | X | X | X |

### 2.3 Neural network architecture

We built regression models using fully connected neural networks to learn and then infer species diversity based on the climatic, geographic, and human footprint features described above. While the values that can be used to train an NN regression model can theoretically take any range, it generally helps the model to converge when rescaling these values to a smaller range, approximately ranging between 0 to 1. We therefore rescaled our training labels by dividing the diversity values by the following scaling divisors, which were approximated to match the maximum values found in the training data for each diversity metric: alpha scaling divisor = 100, beta scaling divisor = 1, gamma scaling divisor = 800.

Models differed in their number of hidden layers and number of nodes per layer (see model testing below, Table 2). Further, we applied different fractions of dropout in our models, which leads to randomly “dropping” the specified fraction of nodes in each hidden layer in each training epoch. This has the effect of reducing overfitting towards the training data, as the model is forced to rely less on individual highly optimized weights. We used the rectified linear units function (ReLU) as the activation function within each layer, and a softplus activation function for the output layer. The softplus activation function in the output layer ensures that the output values (diversity estimates) are all within a positive range, while not imposing any restrictions on the possible maximum value.

**Table 2:** Prediction accuracy for test set of all tested models. The last three columns show the mean average percentage error (MAPE) of the model predictions for an independent test set. The tested models differ in terms of the number of features (first column), number of layers and nodes per layer (second column), and the dropout rate (third column). The best models for each diversity metric are highlighted in bold. More detailed visualizations of the test set predictions for these best models are shown in Fig. 4.

| Features | Nodes | Dropout | Alpha | Beta | Gamma |
| --- | --- | --- | --- | --- | --- |
| 6 | 30 | 0 | 0.6611 | 0.0750 | 0.1088 |
| 6 | 30 | 0.1 | 0.7078 | 0.0773 | 0.1409 |
| 6 | 30 | 0.3 | 0.7095 | 0.0779 | 0.1434 |
| 6 | 30, 5 | 0 | 0.6440 | 0.0752 | 0.1013 |
| 6 | 30, 5 | 0.1 | 0.6129 | 0.0761 | 0.1356 |
| 6 | 30, 5 | 0.3 | 0.7103 | 0.0788 | 0.1457 |
| 6 | 30, 15, 5 | 0 | 0.6570 | 0.0751 | 0.0823 |
| 6 | 30, 15, 5 | 0.1 | 0.6111 | 0.0752 | 0.0951 |
| 6 | 30, 15, 5 | 0.3 | 0.6725 | 0.0783 | 0.1312 |
| 6 | 30, 20, 10, 5 | 0 | 0.6225 | 0.0743 | 0.0804 |
| 6 | 30, 20, 10, 5 | 0.1 | 0.6542 | 0.0749 | 0.1012 |
| 6 | 30, 20, 10, 5 | 0.3 | 0.6844 | 0.0794 | 0.1307 |
| 8 | 30 | 0 | 0.6555 | 0.0742 | 0.1064 |
| 8 | 30 | 0.1 | 0.7022 | 0.0753 | 0.1056 |
| 8 | 30 | 0.3 | 0.6776 | 0.0763 | 0.1107 |
| 8 | 30, 5 | 0 | 0.6301 | 0.0749 | 0.0851 |
| 8 | 30, 5 | 0.1 | **0.5872** | 0.0757 | 0.1012 |
| 8 | 30, 5 | 0.3 | 0.6740 | 0.0779 | 0.1298 |
| 8 | 30, 15, 5 | 0 | 0.6179 | 0.0745 | 0.0673 |
| 8 | 30, 15, 5 | 0.1 | 0.6335 | 0.0749 | 0.0911 |
| 8 | 30, 15, 5 | 0.3 | 0.6606 | 0.0778 | 0.1173 |
| 8 | 30, 20, 10, 5 | 0 | 0.6157 | 0.0741 | 0.0634 |
| 8 | 30, 20, 10, 5 | 0.1 | 0.6047 | 0.0731 | 0.0877 |
| 8 | 30, 20, 10, 5 | 0.3 | 0.7323 | 0.0788 | 0.1357 |
| 27 | 30 | 0 | 0.6233 | 0.0732 | 0.0882 |
| 27 | 30 | 0.1 | 0.6198 | 0.0741 | 0.0829 |
| 27 | 30 | 0.3 | 0.6336 | 0.0750 | 0.0954 |
| 27 | 30, 5 | 0 | 0.6073 | 0.0738 | 0.0835 |
| 27 | 30, 5 | 0.1 | 0.5884 | 0.0736 | 0.0835 |
| 27 | 30, 5 | 0.3 | 0.6157 | 0.0764 | 0.1016 |
| 27 | 30, 15, 5 | 0 | 0.5921 | **0.0721** | **0.0609** |
| 27 | 30, 15, 5 | 0.1 | 0.6165 | 0.0747 | 0.0819 |
| 27 | 30, 15, 5 | 0.3 | 0.6343 | 0.0791 | 0.1145 |
| 27 | 30, 20, 10, 5 | 0 | 0.5904 | 0.0722 | 0.0660 |
| 27 | 30, 20, 10, 5 | 0.1 | 0.6153 | 0.0740 | 0.0987 |
| 27 | 30, 20, 10, 5 | 0.3 | 0.6824 | 0.0786 | 0.1221 |

During training, the network was set to optimize for minimum mean absolute error (MAE) between the (rescaled) true diversity values and the network predictions. Of the 7,896 training instances (vegetation plot sites), we set aside 20% (1,579 instances) as an independent test set. We assigned another 20% (1,579 instances) of the data as a validation set, which we used to determine the optimal number of training epochs that minimizes the validation set MAE, while preventing overfitting towards the training data. All models were trained with the remaining 60% of the data (4,738 instances), using a batch size of 40 instances.

### 2.4 Model testing and evaluation

We tested a range of different training configurations for each diversity metric, specifically testing different combinations of input features, different numbers of hidden layers and nodes per layer, and different dropout fractions (Table 2). Based on the diversity predictions for our independent test set, we calculated the mean absolute percentage error (MAPE) for each model, which differs from the MAE in being a relative error, scaled by the absolute values of the predictions. For each diversity metric we determined the best model configuration by picking the model with the lowest MAPE score.

After identifying the most suitable settings through model testing, we retrained this best model for each diversity metric, using all 7,896 training instances. To avoid overfitting towards the training data, we trained these production models only until the optimal epoch determined during model testing. For each diversity metric we trained an ensemble of 50 models with different random starting seeds, using the best model settings. We averaged the predictions across all these 50 models for each diversity metric, and also calculated the coefficient of variation (standard deviation divided by mean) as a measure of variation of the predicted diversity values, representing uncertainty.

### 2.5 Prediction data

To produce the predictions of alpha, beta, and gamma diversity across Australia we defined a grid with a cell size of 10×10 km and extracted the 27 features for each of the cell centroids. We set the plotsize feature for all points to 500 m^2^ (most common vegetation plot size in training data, Supplementary Fig. S9). Therefore, the predicted alpha diversity values reflect the expected number of plant species to be found in a plot of size 500 m^2^. The radius feature, describing the size of the surrounding area around a point for which gamma diversity is estimated, was set to 5 km, to approximately match the size of the grid cells (10×10 km square).

By adjusting the radius feature, our trained models can be used to predict beta and gamma diversity at user-defined spatial resolutions, as it can be adapted to match the given cell size. Similarly, adjusting the plot size feature allows us to predict alpha diversity for any given plot size. This enables great flexibility in predicting species diversity at different spatial resolutions of the prediction grid, while inherently accounting for species-area relationships, as these are learned by the model. For both, the radius feature as well as the plot size feature, the selected values for prediction should be chosen to be within the range of values present in the training data (Supplementary Fig. S9).

## 3 Results

An overview of all tested models is shown in Table 2. The same model configuration was identified as the best model for beta and gamma diversity: all 27 features, 3 layers with 30, 15, and 5 nodes, respectively, and no dropout (dropout rate = 0). For alpha diversity, on the other hand, we identified as the best model the following configuration: 8 features (see Table 1), 2 layers with 30, and 5 nodes, and a dropout rate of 0.1. We identified the following training epochs as the stopping points for model training, as they constituted the best compromise between optimal model training and avoiding overfitting towards the training set (rounded to the nearest 50): 1500 epochs (alpha), 750 epochs (beta), and 1700 epochs (gamma, see Supplementary Fig. S10). We used these numbers of training epochs for training of the 50 productions models for each diversity metric.

The best alpha diversity model predicted the test set, consisting of approximately 1,600 vegetation plots, with a mean absolute percentage error (MAPE) of 58.72% (Fig. 3). This means that the predicted diversity for the average test set instance was within an approximately 60% range of the true diversity value. This comparably high prediction error is likely caused by the fact that the alpha diversity training instances show a complex spatial pattern, with no easily discernible areas of high or low diversity (Fig. 1). The fact that most of the training features are spatially autocorrelated (such as the BioClim climatic layers) makes it difficult for the model to deduct any meaningful signal from these features during training for predicting alpha diversity. The predictions made by an ensemble of 50 trained alpha models show comparably large uncertainties in some areas (Fig. 4), with a median coefficient of variation across all cells of 0.30. The areas of highest uncertainty – exceeding the median value – are located mostly in the western half of Australia (grey areas in Fig. 4), presumably due to the limited training data from those regions (Fig. 1).

**Figure 3:**
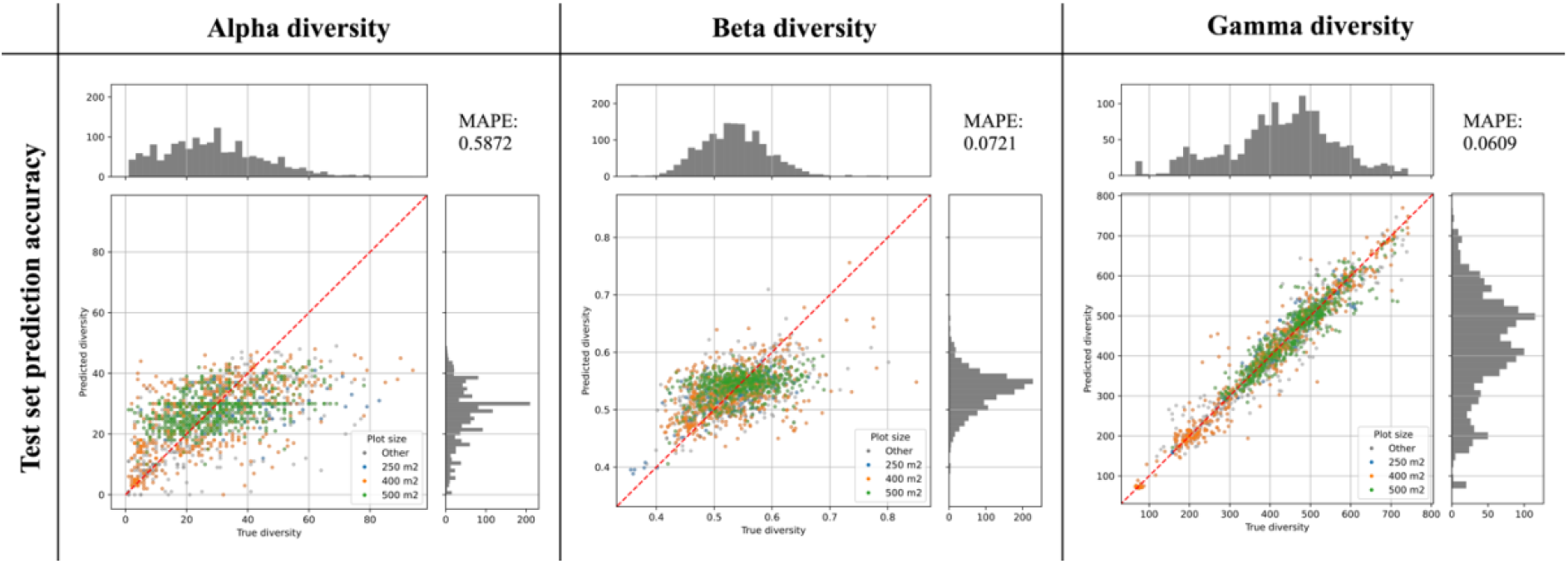
Prediction accuracy of best models as determined on an independent test set. The scatter plots show the predicted diversity (y-axes) plotted against the true diversity (x-axes) for the best alpha, beta, and gamma diversity models. These estimates were made for a randomly selected and independent test set (N = 1,579 instances), exclusively consisting of instances that were not used during model training. The points are colored by the vegetation plot-size associated with each data point (see legend). The red diagonal line shows for reference the best-case scenario, if all labels were predicted 100% accurately. Histograms show the total distribution of values for the true diversity values (top) and the predicted diversity values (right). For each model we calculated the Mean Absolute Percentage Error (MAPE), shown in the top-right corner of each plot.

**Figure 4:**
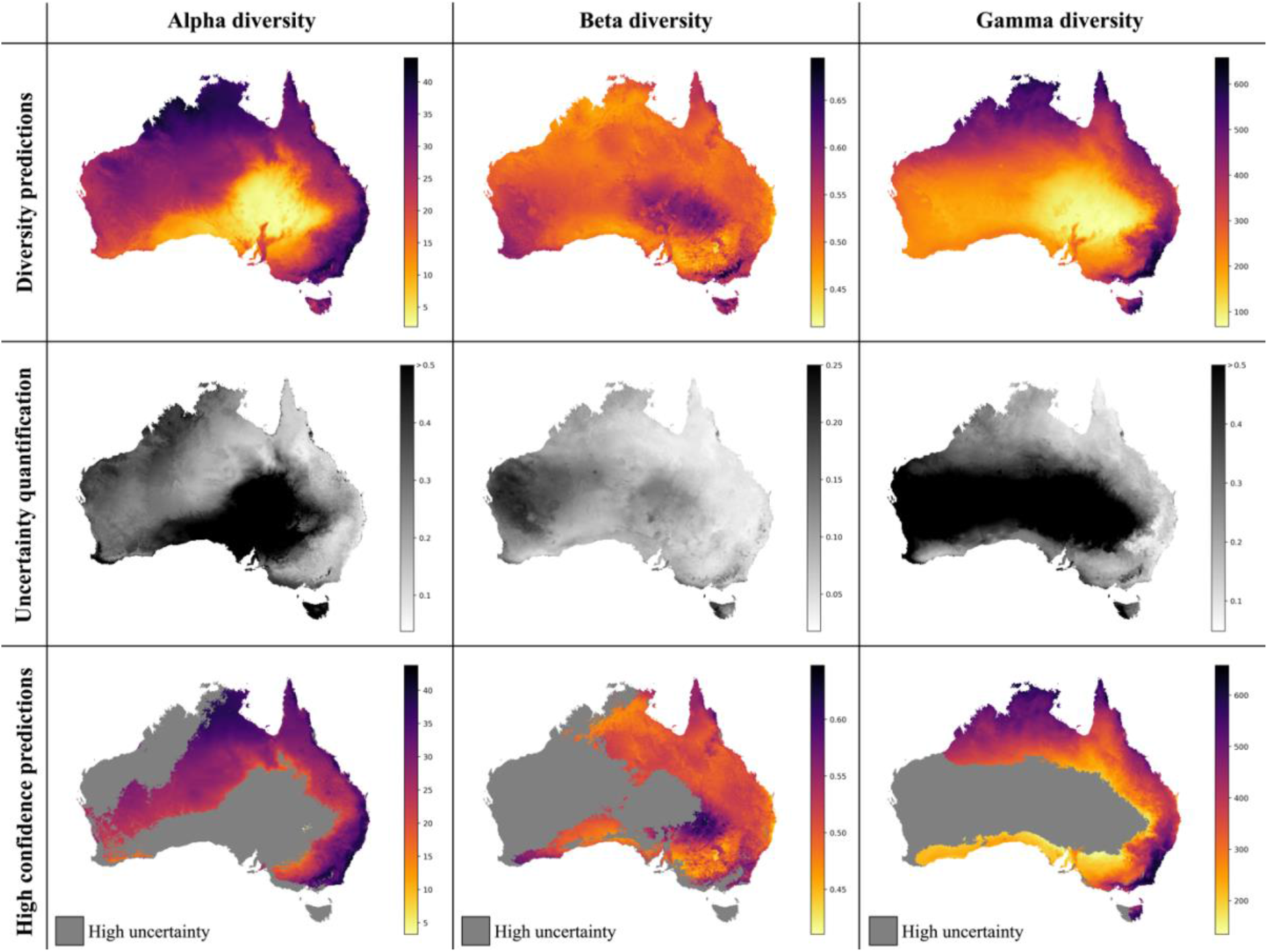
Neural Network predictions for alpha, beta, and gamma diversity of vascular plants. The NN models were trained separately on alpha, beta, or gamma diversity estimates, which we compiled from vegetation plot data (Fig. 1). The alpha diversity maps (left) show the expected number of vascular plant species expected to be found in a 500 m2 plot (most common plot-size found in the vegetation plot data, Supplementary Fig. S2). The beta diversity maps (center) quantifies the spatial turnover and differences in species compositions (Sørensen dissimilarity index, relative to the total diversity) between such 500 m2 plots within each grid cell (10×10 km). The gamma diversity maps show the total species richness within each grid cell. The top row shows the predictions averaged across an ensemble of 50 independently trained models, using different starting seeds. The center row shows the coefficient of variation for each grid cell, as a measure of prediction uncertainty. High values (dark grey/black) correspond to grid cells with less consistent diversity predictions. The bottom row shows the average diversity predictions for only those grid cells with the most consistent diversity predictions (coefficient of variation smaller than median across all grid cells), while high-uncertainty grid cells are marked in grey.

The overall highest alpha diversity predictions are found along the eastern coast of Australia, from the northernmost tip of Queensland to the most southwestern part of Victoria (Fig. 4). A potential drop in alpha diversity is visible in the area around Cairns, extending about 100 km south from the city area, perhaps corresponding with the Burdekin-Lynd gap (Edwards et al., 2017), yet these grid cells are predicted with comparably high uncertainty, giving only weak support for this observed pattern. Other areas of medium to high alpha diversity inferred by our model are the top end of the Northern Territory, as well as the north Kimberley in northern Western Australia.

The best beta diversity model showed an MAPE score of 7.21%, constituting a substantially higher accuracy compared to the alpha diversity model. Similarly, the median coefficient of variation across all prediction grid cells was very low with 0.09, indicating high consistency in the predicted diversity pattern. The high-uncertainty cells, identified as having a coefficient of variation above the median, largely overlap with those identified for the alpha diversity model, covering the majority of Western Australia (Fig. 4). Perhaps being the least intuitive of the three diversity metrics, areas with a high predicted beta diversity within our framework represent sites that are expected to show large differences in species composition between vegetation plots within the defined area (a given grid cell).

Differently to alpha diversity, the majority of the eastern coastal areas show medium to low beta diversity values. Higher beta diversity is inferred for the southeastern part of Australia, particularly in higher elevations between Canberra and Melbourne. High species turnover is also inferred for the arid eastern desert of central Australia, as well as for south-western Australia.

With a MAPE score of only 6.09%, our gamma model performed the best out of the three different diversity metrics prediction models. The median coefficient of variation of gamma predictions across all of Australia was quantified at 0.37. As for the other two models, this variation was largely driven by high uncertainty grid cells in the western half of the continent (Fig. 4). Below we discuss the specific spatial diversity patterns present that were predicted by our models in more detail (see Discussion).

When evaluating our model predictions on a per-biome basis, excluding high uncertainty predictions as identified in Fig. 4, we identify differences in predicted diversity between biome types (Fig. 5). For alpha and gamma diversity, we find the highest average diversity predictions for tropical forests, temperate forests, montane shrublands and grasslands, and tropical and subtropical grasslands and savannas. Our beta diversity estimates, on the other hand, show a rather uniform pattern across biomes, with the exception of montane grasslands and shrublands, which show the highest species turnover. The high beta diversity identified for the montane biome may be driven by the increased elevational gradients in this area, as species turnover has been found to be higher along elevational gradients (Venn et al., 2017; Albrecht et al., 2021).

**Figure 5:**
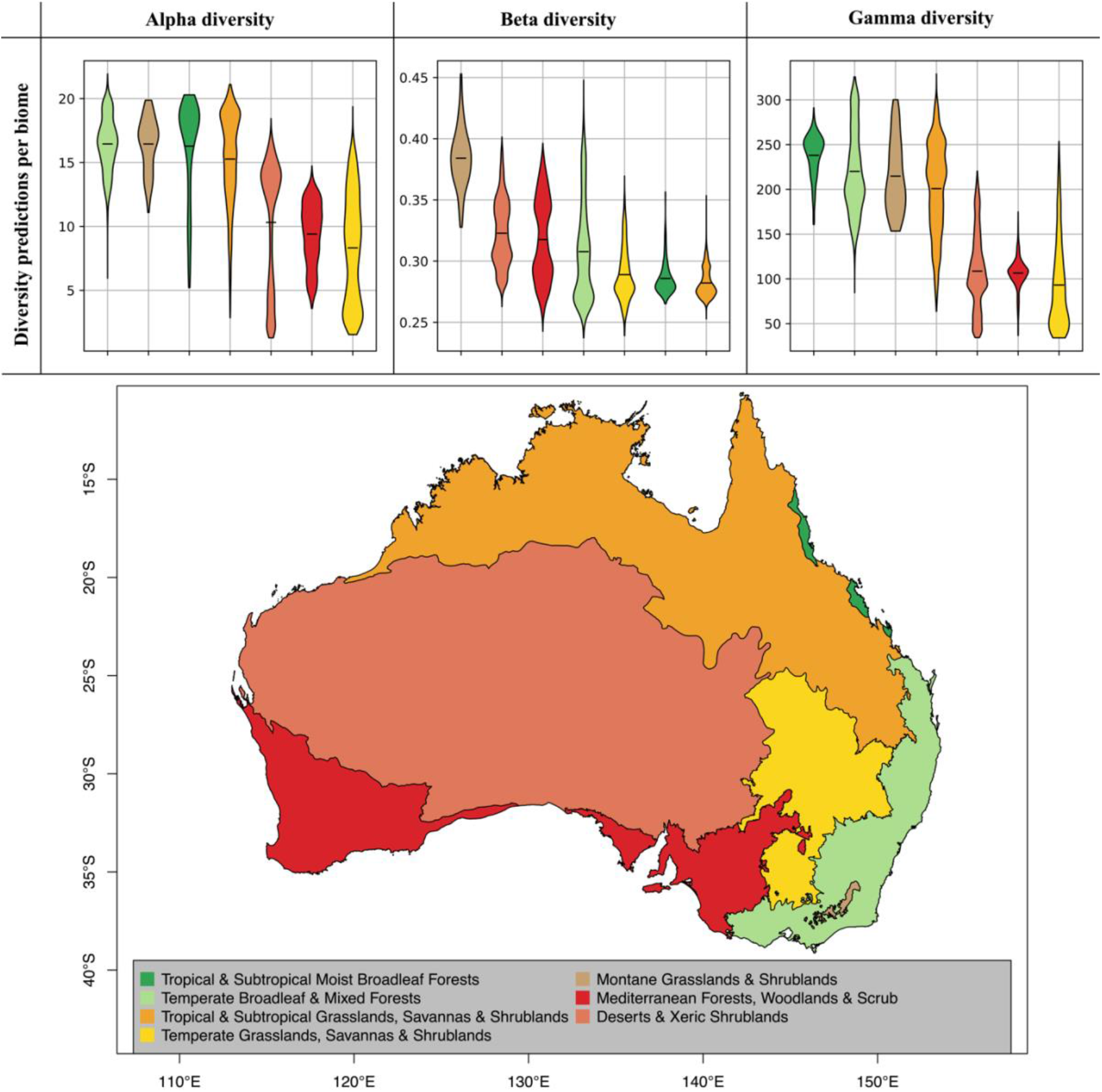
Diversity predictions by biome. The violine plots show the range of diversity predictions across all grid cells within a given biome, excluding high uncertainty predictions (see. Fig. 4). The horizontal black lines inside the violine plots mark the mean estimate for each biome. The biomes, which are displayed on the map, were compiled from the Terrestrial Ecoregions of the World (TEOW) data (Olson et al., 2001).

## 4 Discussion

### 4.1 Using neural networks for diversity predictions

Here we developed and applied a novel approach of estimating species diversity, using neural networks. We showcased our model, using vegetation plot data that is openly available through the sPlotOpen database for Australia, and showed that it can be used to accurately predict diversity on different scales (alpha, beta, and gamma) for any given point in Australia. This enables us to produce maps of species diversity at a wide range of spatial resolutions. The novelty of our approach, as compared to previous approaches of modelling species diversity, is that i) it does not require the modelling of distribution ranges for individual species (Mutke and Barthlott, 2005; Barthlott et al., 2007), ii) it does not require an *a priori* definition of species-area relationships (Kier et al., 2005), iii) it does not require the assumption of monotonic and usually oversimplifying relationships (e.g. linear or exponential) between predictors and response variable (Cingolani et al., 2010), and iv) it allows the direct quantification of uncertainty in the predictions.

Given these advantages, and the easy combination of different features of continuous or categorical nature, our deep learning model, represents a promising new tool for the task of predicting diversity. This study and other recent work (e.g., Večeřa et al., 2019) demonstrate how such models can be trained on readily available data from public databases. The accuracy of these models could be potentially further improved by compiling additional features deemed to be informative for the task of diversity prediction. Preferably such features should be based on data available in form of a spatial grids covering the entirety of the prediction area (in this case, Australia). Remote sensing data are a promising and potentially highly informative data source to fill up spatial gaps with increasingly detailed vegetation maps (Gholizadeh et al., 2020; Moat et al., 2021), and could be applied in future machine learning models for the task of diversity prediction.

### 4.2 Correlation between diversity metrics

Previous studies have found all three diversity metrics to be correlated (Cingolani et al., 2010). Here we find that the maps produced for alpha and for gamma diversity overall show similar diversity hotspots, while beta diversity shows a different spatial pattern (Figs. 4 and 5). There is a wide variety of definitions of beta diversity, some which are directly correlated to alpha and gamma diversity (e.g., Whittaker’s original definition of *β* = *γ/α*, sensu Whittaker, 1960). However, the Sørensen dissimilarity index *β_sor_* used in this study does not display such a direct correlation to either alpha or gamma diversity, leading to the distinctly different spatial pattern observed in our predictions, which reflects the different patterns between this metric and the other two also observed in the training data (Fig. 1).

While the patterns of alpha and gamma diversity inferred by our models are strongly correlated, they do differ in some areas. There is potential for areas with low gamma diversity to exhibit relatively high densities of species, leading to high alpha diversity estimates within smaller defined areas, such as the 500 m2 vegetation plots predicted by our models. This is particularly the case for vegetation types consisting of species with relatively small individual plant sizes (such as grasslands and shrublands), which in comparison with forests allow for a potentially denser accumulation of individuals. These differences in average plant size often lead to open habitat grasslands displaying comparatively high alpha diversity values, particularly on small plot sizes (Wilson et al., 2012).

### 4.3 Spatial biases in training data

Sampling biases pose a severe challenge for biodiversity reconstruction in countries of uneven spatial sampling, such as Australia (Piccolo et al., 2020). In our approach, we account for geographic bias in the training data by quantifying the uncertainty in the diversity predictions, which largely reflect those areas with little or no training instances. Additionally, we add the count of GBIF occurrence records in the surrounding of any given training instance as a measure of general sampling effort. Recent studies have addressed the issue of differences in sampling effort in more detail for defined regions and have pointed a way forward in addressing and accounting for this issue, using strategically sampled empirical data (Gioia and Hopper, 2017). However, such efforts are labor- and time-intensive and may not be feasible on continental scales. Alternatively, computational tools that can readily quantify spatial biases based on public database data are a promising way forward towards better accounting for the issue of spatial sampling biases (Zizka et al., 2021).

### 4.4 Predicted diversity patterns for Australia

Our model predictions of alpha and gamma diversity identify several vascular plant biodiversity hotspots for Australia, such as i) the tropical and subtropical forests in northeastern Queensland, ii) the temperate forests and the montane grasslands and shrublands of southeastern Australia, iii) the tropical savanna dominated ecosystems of the Northern Territory, and iv) northern Western Australia (Figs. 4 and 5). These areas of high vascular plant diversity largely correlate with findings of previous studies, e.g. (Steffen, 2009; Goldie et al., 2010; Yeates et al., 2014; Thornhill et al., 2016) and are highly correlated with broader climatic patterns (Ooi et al., 2017).

One notable difference of our model predictions compared to previous work is the south-west of Western Australia, which is often inferred as a plant diversity hotspot (e.g. Myers et al., 2000; Steffen, 2009), but was predicted with comparably low alpha and gamma diversity by our models. This south-west Australian biodiversity hotspot may not have been predicted accurately – as also indicated by the large prediction uncertainty identified by our model – due to alternate evolutionary patterns in the region that have led to higher diversity than might otherwise be predicted in this very old and climatically buffered, infertile landscape (an OCBIL; see Hopper et al., 2016). It is also interesting to note that the models predict similar alpha diversity between the Kimberley region of Western Australia and the top end of the Northern Territory, as recent surveys demonstrate that this is indeed the case (R.L. Barrett & M.D. Barrett, unpubl. data).

Interestingly, our beta diversity model inferred high species turnover for the arid eastern desert of central Australia. While this region has the lowest estimates for alpha and gamma diversity, the species turnover (relative to the total diversity) is inferred to be among the highest on the continent, likely reflecting a complex mosaic of Mediterranean, temperate and arid vegetation types in this region (Fox, 2007).

## 6 Author Contributions

TA, AA, and DS contributed to conception and design of the study. TA compiled the data, wrote the code, and ran all analyses. TA wrote the first draft of the manuscript, with contributions of sections written by AA, RB, and DS. All authors contributed to manuscript revision, read, and approved the submitted version.

## 8 Acknowledgments

We thank the members of the Gothenburg Global Biodiversity Centre (GGBC) for valuable feedback during the early stages of this project, in particular Søren Faurby.

## 9 Supplementary Material

The supplementary material accompanying this study contains Supplementary Figures S1-S13.

## 10 Data Availability Statement

Publicly available datasets were analyzed in this study. These data can be found here: **https://doi.org/10.5281/zenodo.5792187**. All code developed here (Python and R scripts) will be made available here: **https://github.com/tobiashofmann88/plant_div_NN**.

