## Supplementary Figures S1-S13 for "Estimating alpha, beta, and gamma diversity through deep learning"

### *Supplementary Material*

#### 1 Supplementary Figures

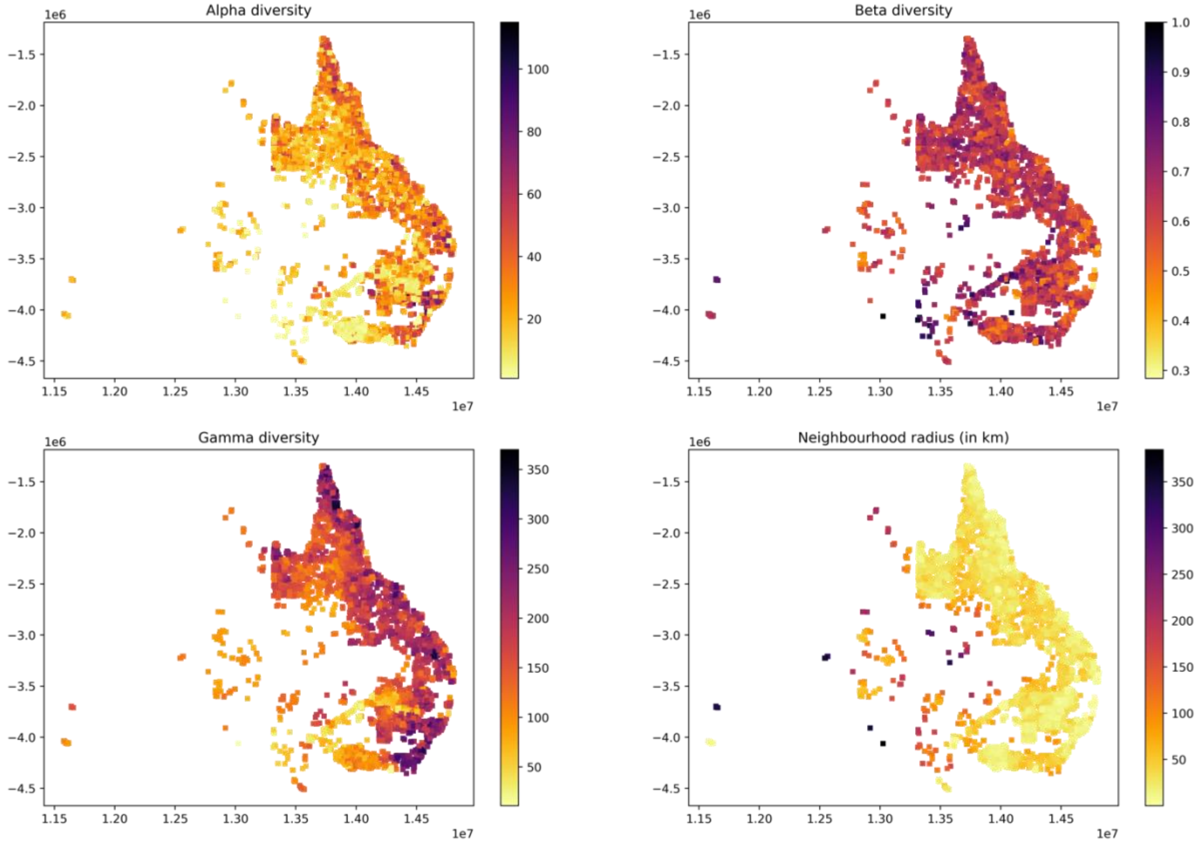


**Supplementary Figure S1**: Distribution of vegetation plots and diversity measures used for training for N=10. The panels show the different diversity measures that were used for model training (see panel titles). The last panel shows the radius of the area containing the N nearest neighbours of each point, which was used for the calculation of beta and gamma diversity.

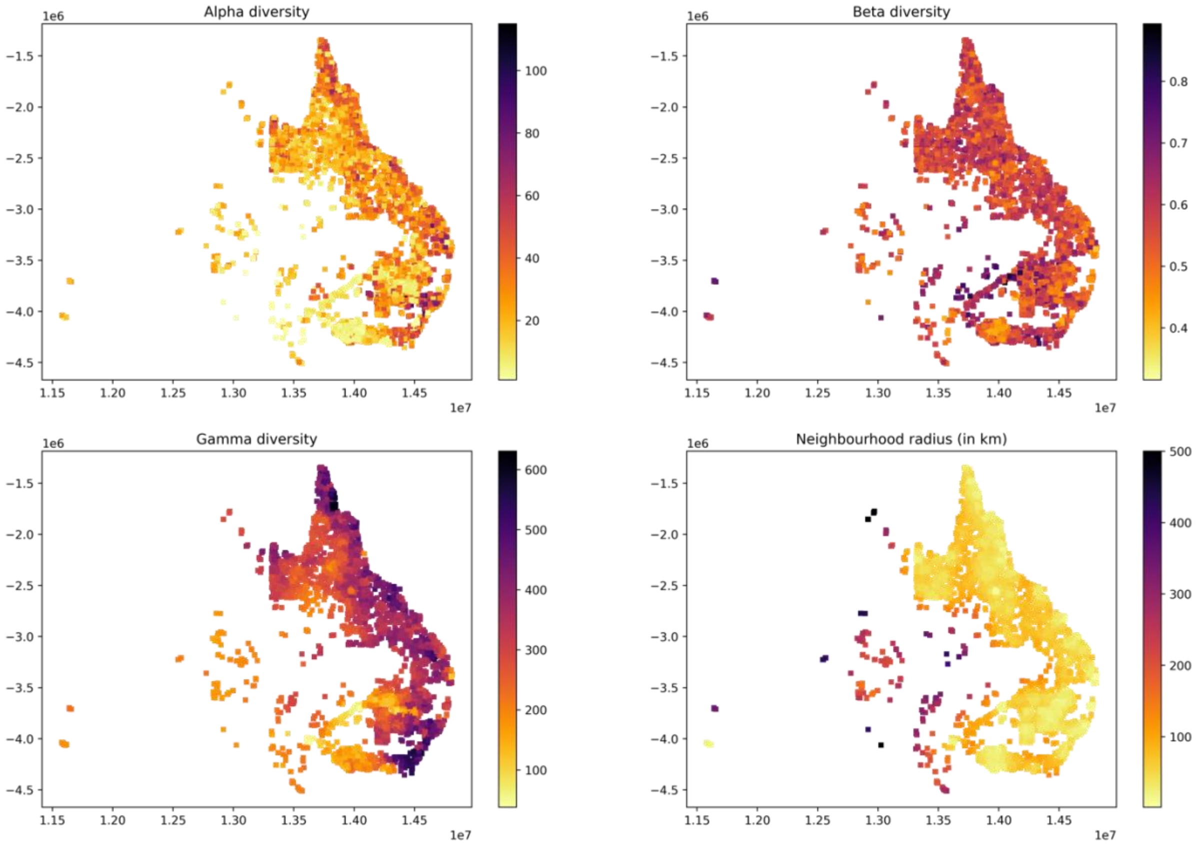


**Supplementary Figure S2**: Distribution of vegetation plots and diversity measures used for training for N=30.

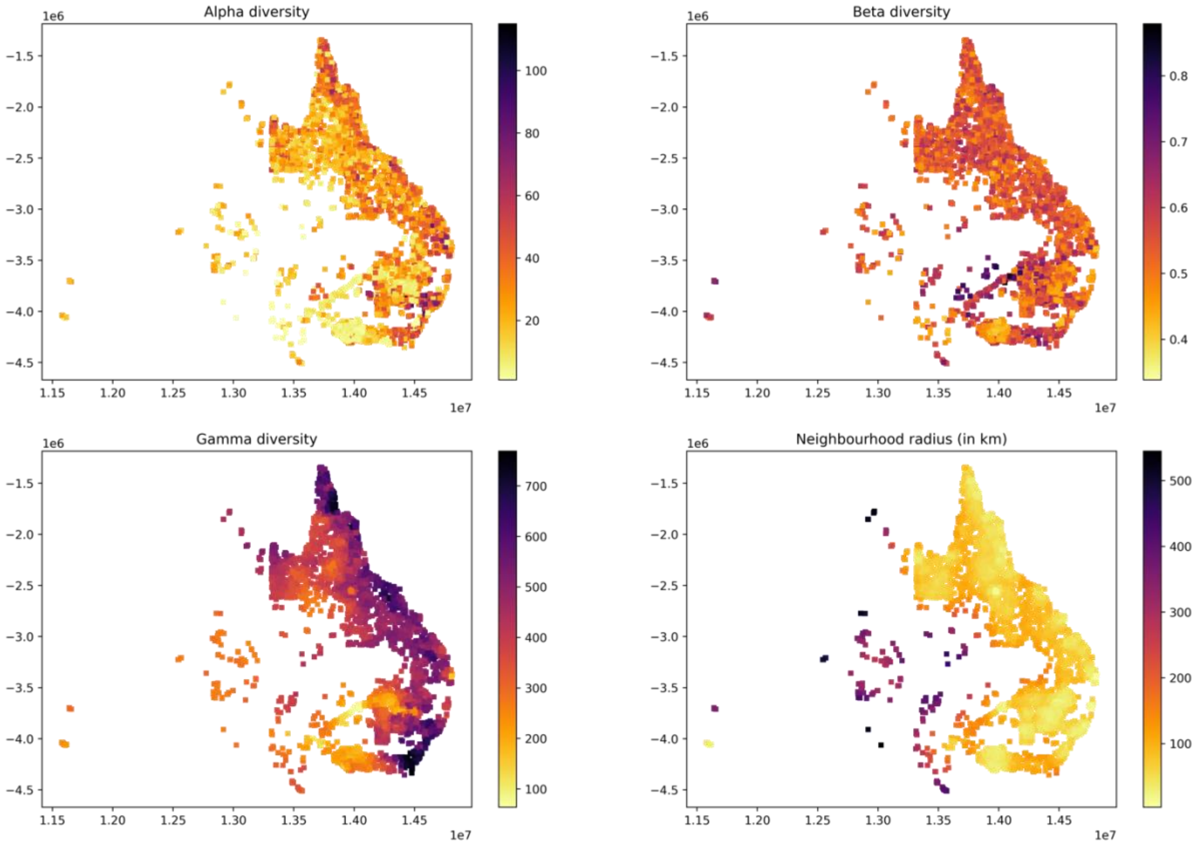


**Supplementary Figure S3**: Distribution of vegetation plots and diversity measures used for training for N=50.

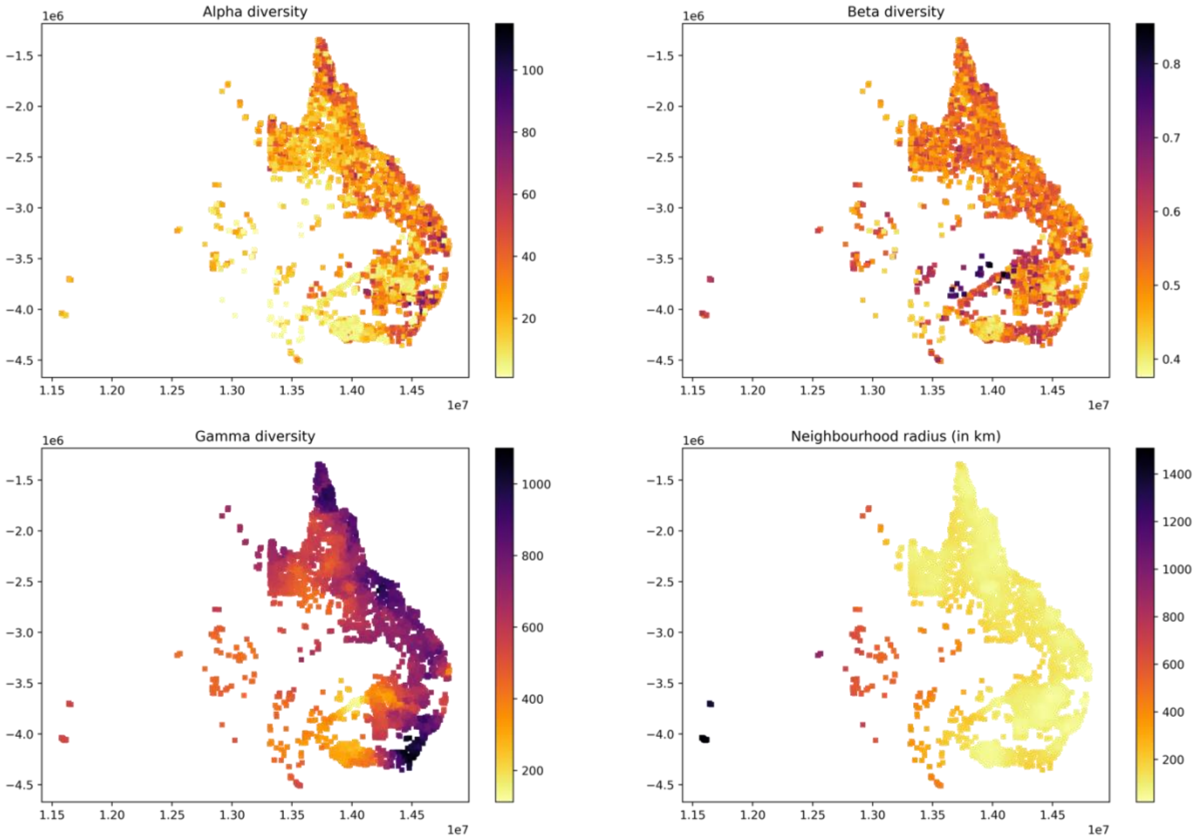


**Supplementary Figure S4**: Distribution of vegetation plots and diversity measures used for training for N=100.

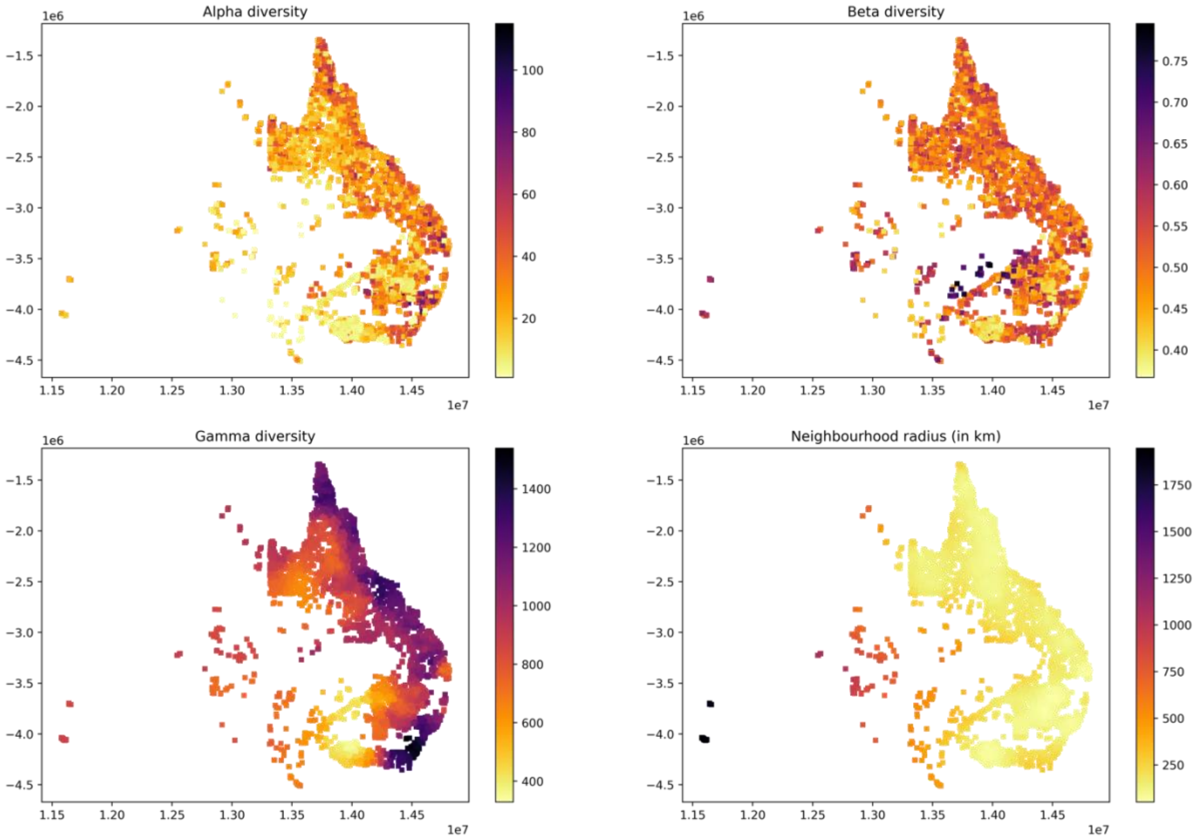


**Supplementary Figure S5**: Distribution of vegetation plots and diversity measures used for training for N=200.

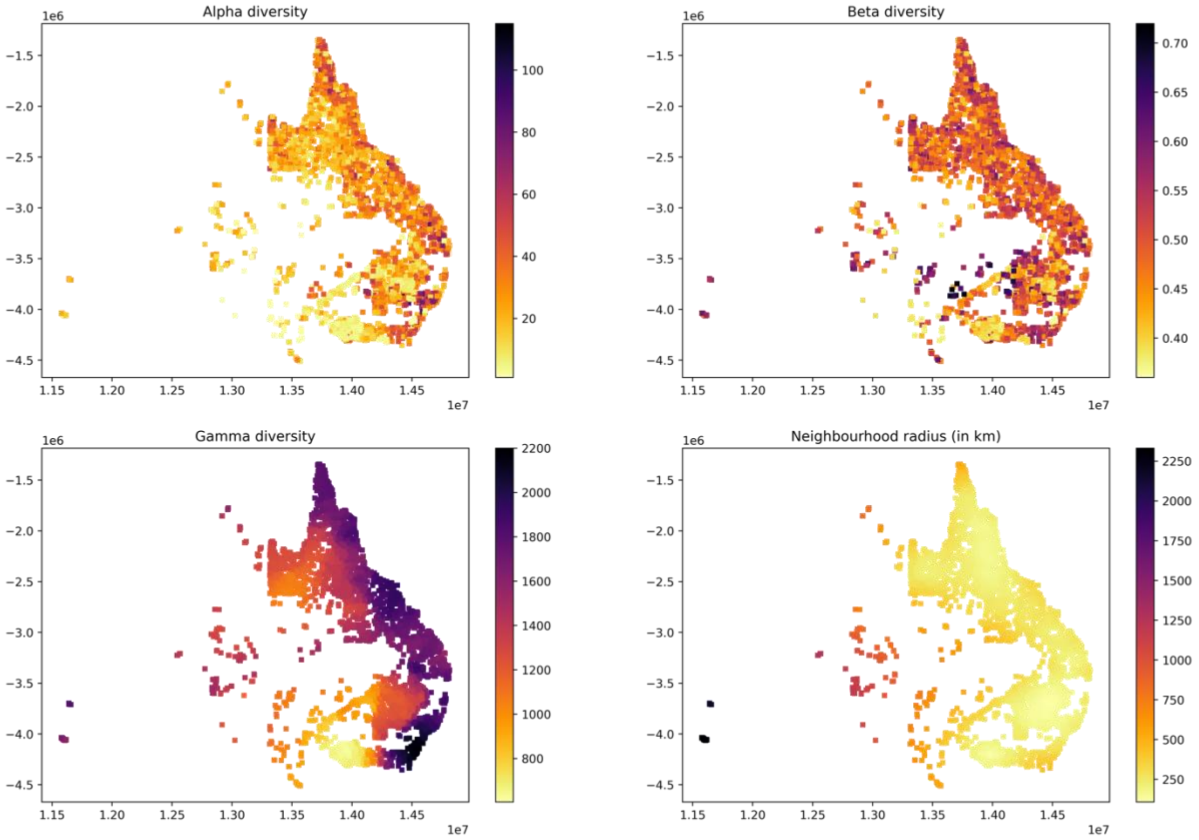


**Supplementary Figure S6**: Distribution of vegetation plots and diversity measures used for training for N=500.

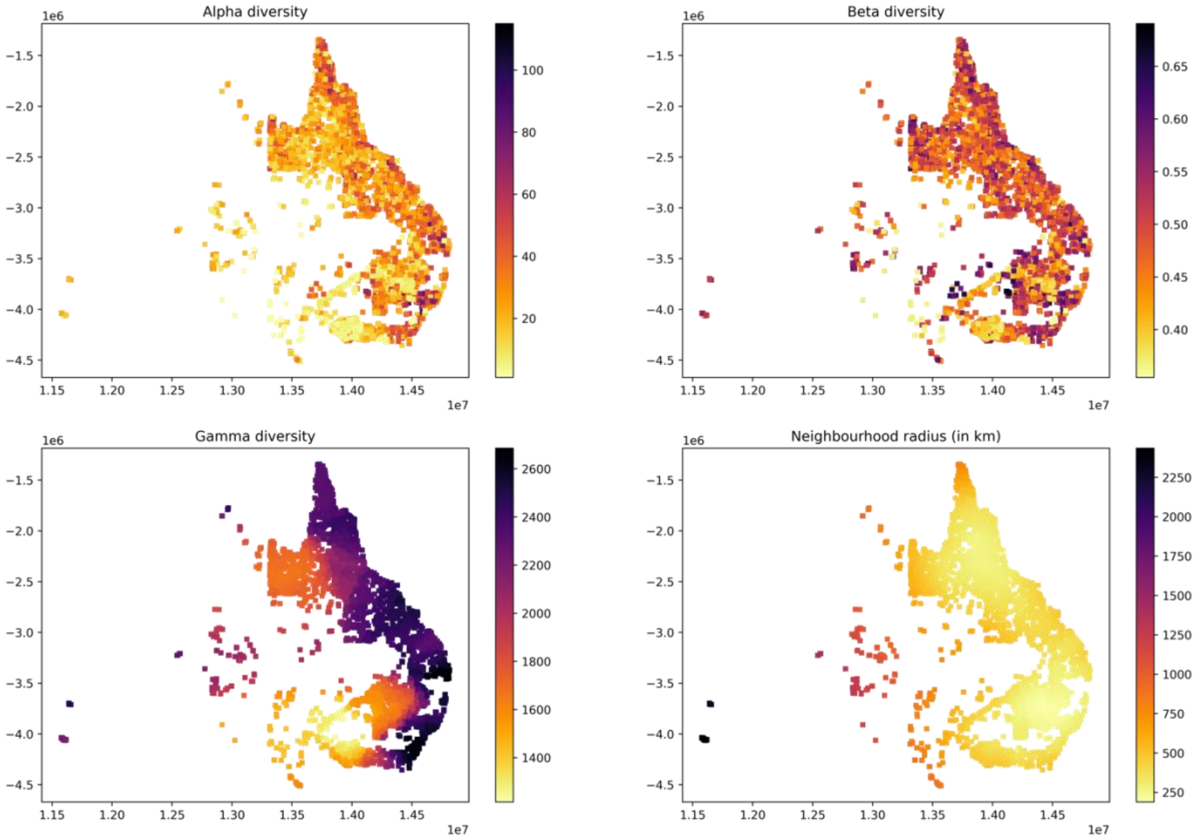


**Supplementary Figure S7**: Distribution of vegetation plots and diversity measures used for training for N=1000.

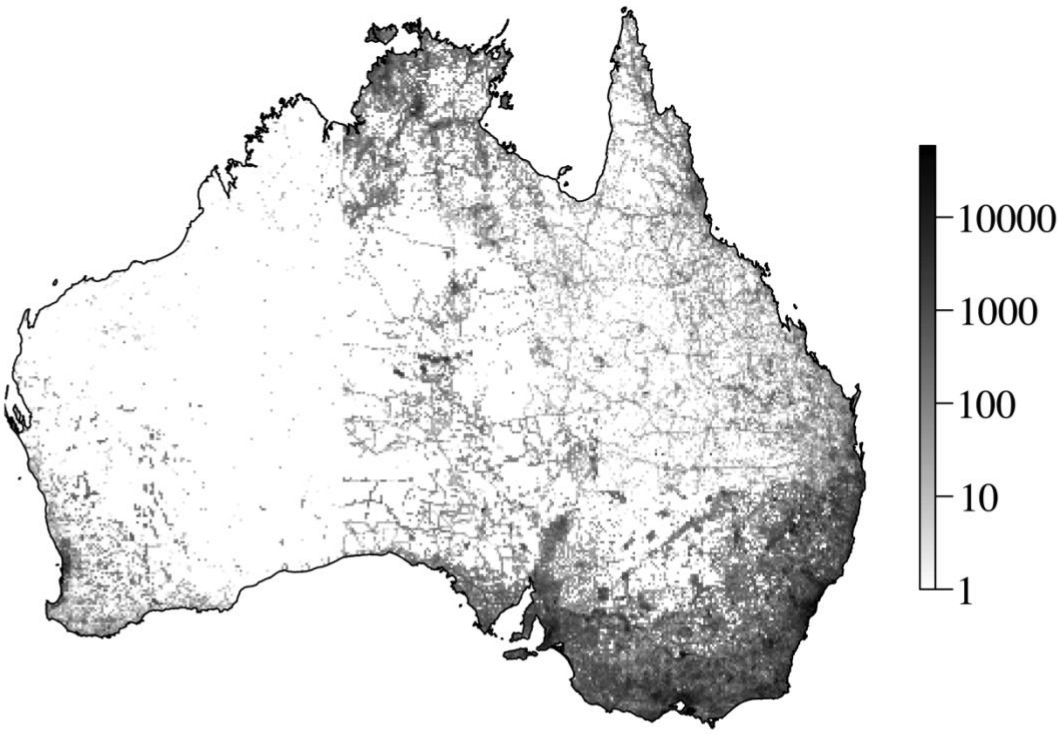


**Supplementary Figure S8: Sampling effort in Australia.** The map shows a 10x10km grid across mainland Australia. The shade of grey corresponds to the counted number of occurrences of vascular plants available through GBIF for each grid cell. Note that the greyscale was applied on a logarithmic basis for better visibility of differences between smaller count values.

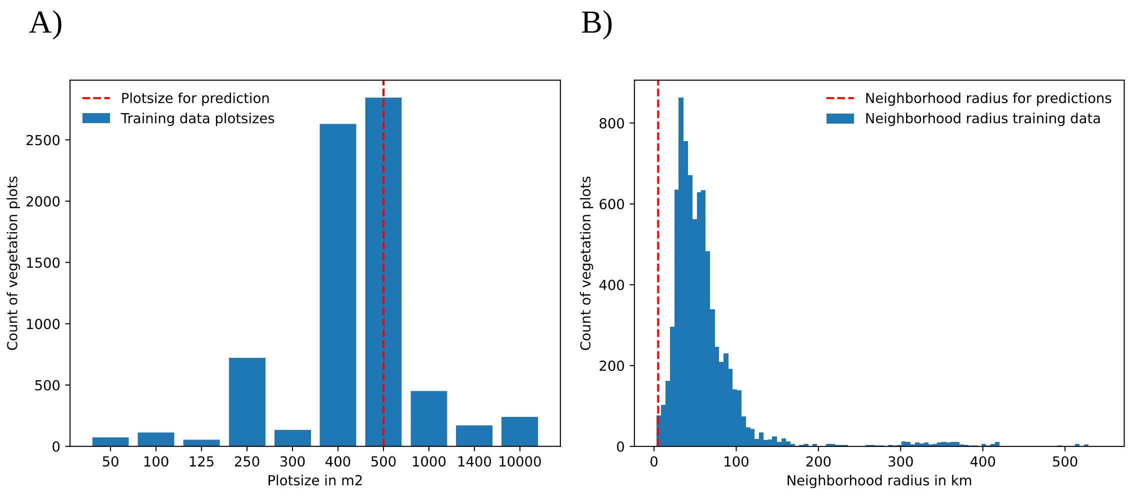


**Supplementary Figure S9: Distribution of adjustable distance features in training data.** The plots show the distribution of vegetation plot sizes (A) and the distribution of neighborhood radiuses, encompassing the 50 nearest neighbors (B) for all training instances.

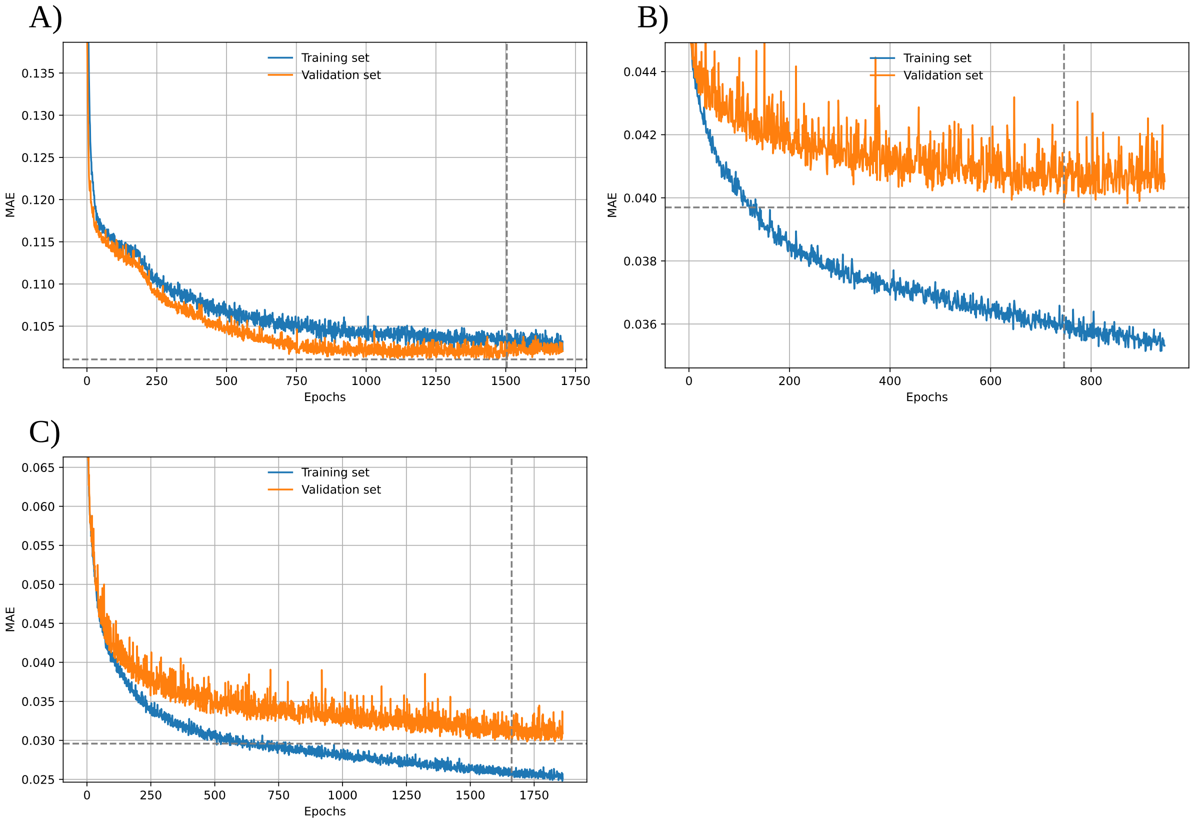


**Supplementary Fig. S10**: Mean absolute error of diversity predictions for training (blue) and validation sets (orange) throughout the training epochs, shown for the best alpha model (A), the best beta model (B), and the best gamma model (C, Table 2). Training was terminated if validation MAE did not improve for 200 consecutive epochs (patience = 200). This termination point (best epochs) is marked by the vertical dashed lines, while the horizontal dashed lines show the MAE value at this best epoch. The area shaded in grey represents the interval of 200 epochs during which no improvement of validation MAE was observed (patience). These last epochs were discarded from model training to reduce overfitting towards the training set.

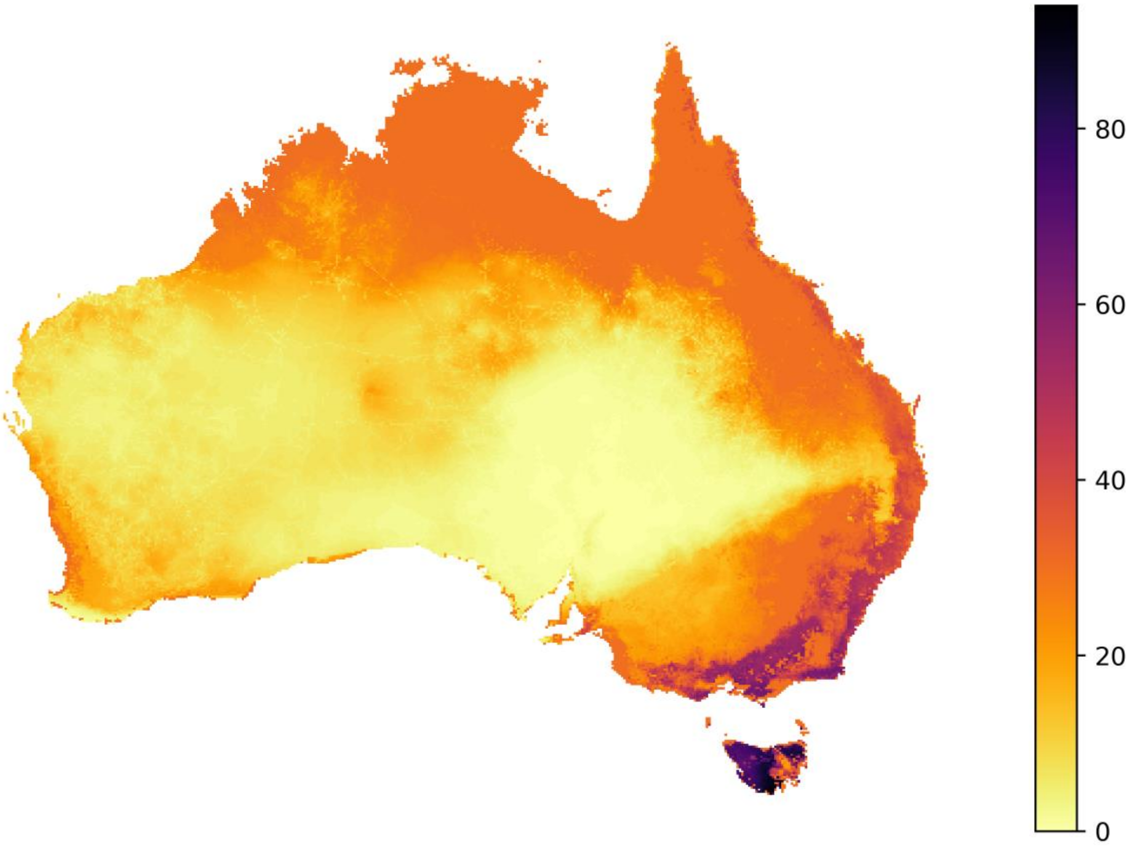


**Supplementary Figure S11**: Alpha diversity predictions of best alpha model. The model was trained using 8 features (Table 1), two hidden layers with 30 and 5 nodes, respectively, and a dropout rate of 0.1. This model resulted in the lowest MAPE score among all tested alpha diversity models during model testing (Table 2). In contrast to the maps displayed in Fig. 4, which constitute the predictions averaged across 50 independently trained models, this map instead shows the predictions of one single model, which are more vulnerable to stochasticity and thus less reliable.

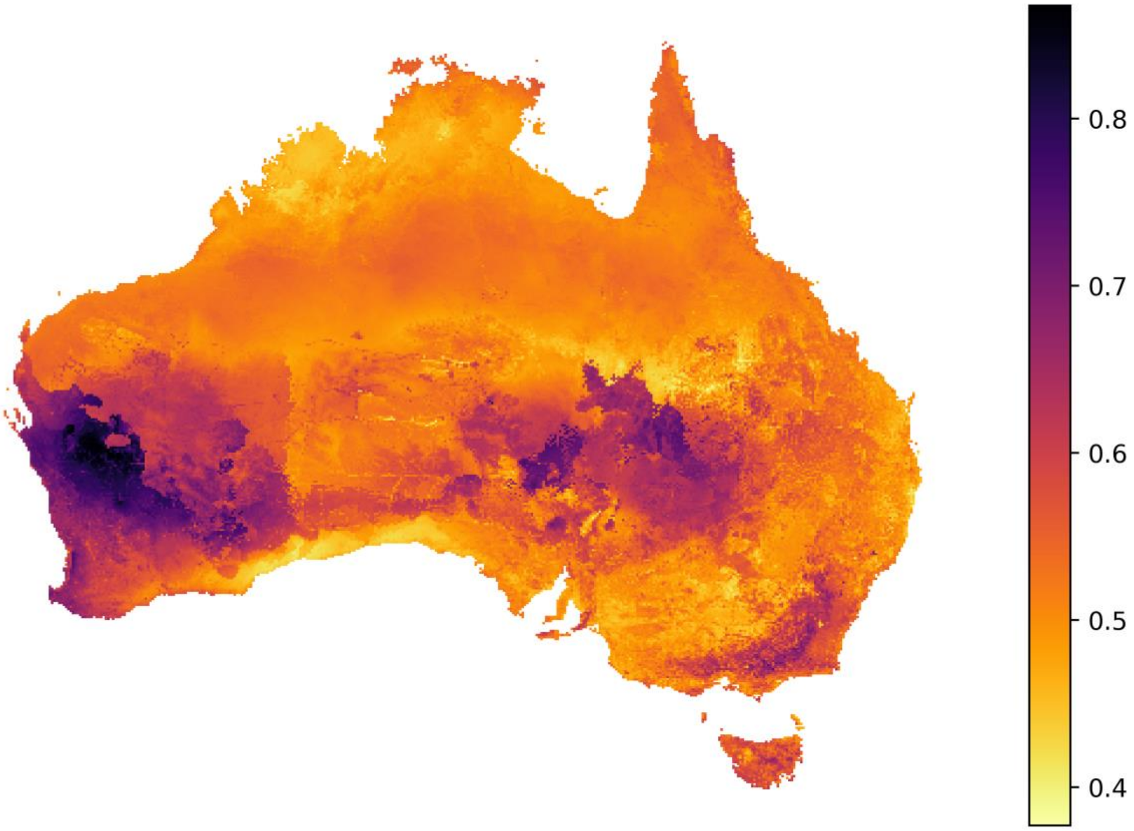


**Supplementary Figure S12**: Beta diversity predictions of best beta model. The model was trained using all 27 features (Table 1), three hidden layers with 30, 15, and 5 nodes, respectively, and a dropout rate of 0 (no dropout). This model resulted in the lowest MAPE score among all tested beta diversity models during model testing (Table 2). In contrast to the maps displayed in Fig. 4, which constitute the predictions averaged across 50 independently trained models, this map instead shows the predictions of one single model, which are more vulnerable to stochasticity and thus less reliable.

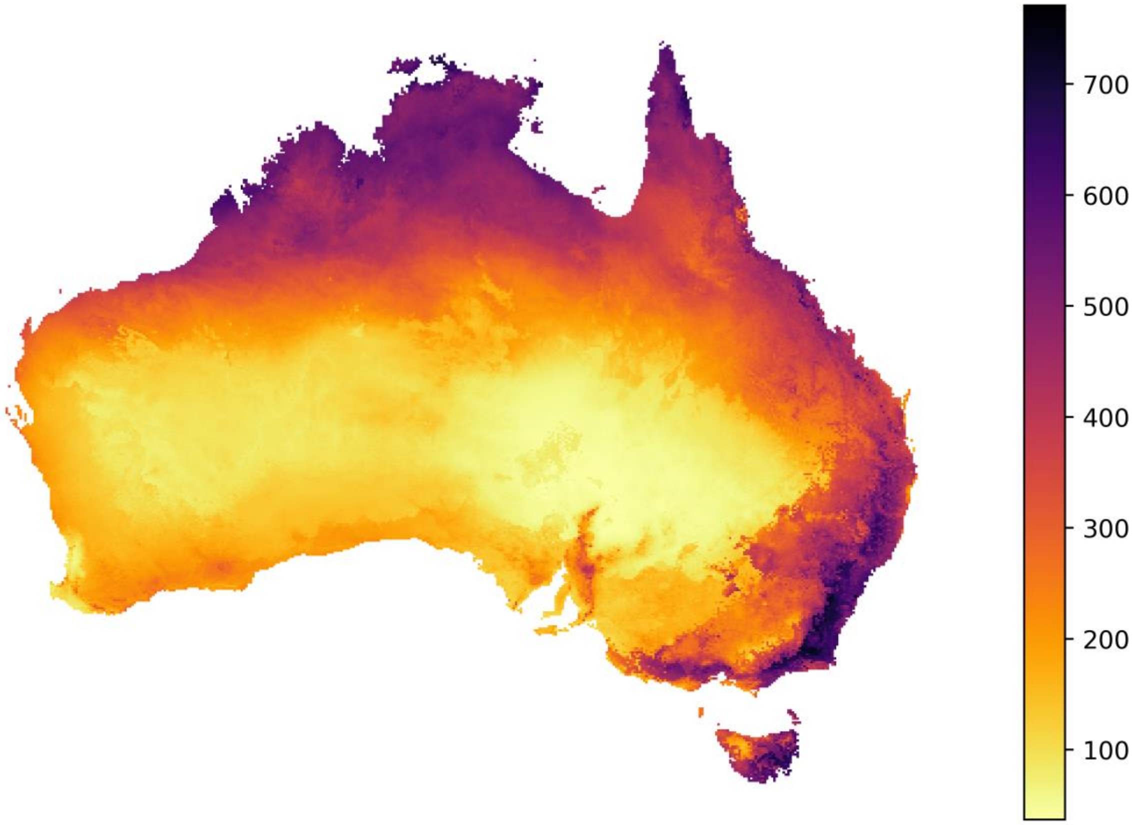


**Supplementary Figure S13**: Gamma diversity predictions of best gamma model. The model was trained using all 27 features (Table 1), three hidden layers with 30, 15, and 5 nodes, respectively, and a dropout rate of 0 (no dropout). This model resulted in the lowest MAPE score among all tested gamma diversity models during model testing (Table 2). In contrast to the maps displayed in Fig. 4, which constitute the predictions averaged across 50 independently trained models, this map instead shows the predictions of one single model, which are more vulnerable to stochasticity and thus less reliable.
